# Expanding the capillarics toolbox: 3D-printed microfluidic phaseguides and self-coalescence modules

**DOI:** 10.1101/2024.12.21.629531

**Authors:** Cosette Craig, Kelsey Leong, Carrie Lin, Megan Chang, Ayokunle Olanrewaju

## Abstract

Capillarics are microfluidic circuits that are assembled from individual fluidic elements, powered by surface tension forces encoded by microchannel geometry and surface chemistry, and enable instrument-free pre-programmed automation of multi-step liquid handling processes. 3D printing has recently transformed capillarics by enabling rapid and cost-effective prototyping, provided addition geometric degrees of freedom in multi-level fabrication, and facilitated new design paradigms with greater capabilities than traditional cleanroom fabrication. Despite widespread interest in 3D printing and development of custom high-resolution stereolithography printers for microfluidic applications, fluidic elements that require precise and tunable control over capillary pinning lines – such as fluidic phaseguides and self-coalescence modules (SCMs) – have so far only been manufactured with centralized and expensive cleanroom methods. Not only does this limit access to versatile capillaric features to only well-resourced settings, but it also slows innovation and widespread application of these fluid handling technologies. Here we expand the toolbox of 3D-printed capillarics to include phaseguides and SCMs, demonstrating their potential for precise instrument-free control over multi-step liquid handling and reagent rehydration. We employed benchtop stereolithography printers to prototype (up to 50X) scaled-up phaseguides and SCMs and integrated them into a capillaric circuit for inline reagent reconstitution, dynamic fluid control, and sequential drainage. We showcased scalable designs, customizable geometries, and robust self-coalescing flow for larger liquid volumes – up to 50 µL compared with 1.25 µL in cleanroom-fabricated SCMs. This work represents a significant advance in democratizing access to microfluidics, with potential for broad applications in diagnostics, assay automation, and organ-on-chip systems.

## Introduction

Capillary microfluidics manipulate fluids by manipulating surface tension effects that are defined by the geometry and surface chemistry of microchannels, thereby removing the reliance on peripheral equipment for pumping and valving^1^. There are a variety of capillary microfluidic flow control elements including stop valves^2^, trigger valves^3^, retention burst valves^4^, capillary pumps^5^, domino valves^1^, phaseguides^6^, and self-coalescence modules^7^ that provide a variety of self-powered and self-regulated fluidic operations. The field of capillary microfluidics is moving towards *capillaric* circuits – inspired by electric circuits and analogies – that consist of assemblies of capillary microfluidic elements that when combined are capable of instrument-free and user-friendly automation of complex multi-step liquid handling processes. Applications and recent demonstrations include the automation of enzyme linked immunoassays (ELISAs) for rapid detection of bacteria in large (∼100 µL) volumes of urine^8^, a thrombin generation assay for continuous subsampling and analysis of plasma coagulation^9^, and pre-programmed sequential delivery of 300 liquids^1^.

3D-printing has transformed our approach to rapid prototyping of capillarics by reducing manufacturing cost and time and enabling new design paradigms with feature sizes ranging from tens to hundreds of microns^4^. Furthermore, capillarics have also recently been prototyped with inexpensive (<$300 USD) liquid crystal display (LCD) stereolithography (SLA) 3D-printers^10,11^, reducing the barrier to entry for researchers and citizen scientists. Despite these advancements in the 3D-printing of capillarics, key fluidic elements that rely on precise and tunable manipulation of capillary pinning lines (CPLs) – like phaseguides and self-coalescence modules (SCMs) – have so far been primarily prototyped with traditional cleanroom fabrication. This is because features like phaseguides and SCMs are traditionally thought to require high resolution features (down to 5 µm wide)^7^ with very smooth surfaces and perpendicular walls to prevent disruption of the CPL, that are not readily achievable using commercially available benchtop 3D printers.

Phaseguides offer precise control over liquid advancement by strategically altering capillary pressure, thereby tackling the crucial challenges of priming behavior and sample retrieval in intricate channel networks^6^. Rails patterned onto substrates dictate the cross-sectional air-fluid interface to control fluid direction, overflow, and filling/emptying patterns. After their initial demonstration in 2011, phaseguide-based microfluidic systems have become widely used, with applications in 3D cell culturing for organ-on-a-chip (OoC) systems ^12–24^, rapid point-of-care diagnostics^25–29^, RNA extraction^30,31^, electroextraction^32^, and magnetic bead-based assays^33^. Outside their utility in controlling liquid priming, phaseguides have also been applied for passive valving and execution of elaborate pre-programmed liquid routing^34^.

SCMs contain a patterned phaseguide-like feature in a reservoir midplane that directs flow to partially fill a space, then fold in on itself, or self-coalesce^7^. When lyophilized reagents are spotted into an SCM, this characteristic doubling-back of flow reconstitutes dry reagents in highly tunable pulses without shear-induced accumulation of reagents at the fluid front. SCMs have been used to automate isothermal DNA amplification assays^7^, bead-based immunoassays for troponin I detection in human serum analogous to lateral flow assays (LFAs),^35^ and enzymatic detection of glucose-6-phosphate dehydrogenase (G6PD)^36^. These applications leverage the precision, modularity, and multiplexing made available by capillarics to integrate precise sample and reagent delivery. Although recent work sought to transfer SCMs to more rapid and accessible manufacturing with inexpensive laser-cut ‘sticker’ microfluidics for reagent rehydration,^37^ that approach required multiple manual steps for device assembly and did not integrate the CPL with additional fluidic elements like vents and stop valves, that often require high-precision fabrication, thereby precluding integration within capillaric circuits and downstream applications.

Here we expand the portfolio of 3D-printed capillarics to include phaseguides and SCMs. We leveraged the degrees of freedom in microchannel geometry and multi-level channel heights offered by 3D-printing to demonstrate robust and functional 3D-printed phaseguides and SCMs scaled up to 50X compared with conventional cleanroom fabrication. We demonstrated a capillaric circuit with an integrated SCM, showcasing the ability to control capillary pinning lines and automate complex tasks including dilution and pre-programmed multi-step liquid delivery, without external instrumentation.

## Experimental

### Materials

Clear Microfluidics Resin V7.0a (CADworks3D, Toronto, ON, Canada) was used to optimize capillary pinning line features and Water-Wash Resin+ (AnyCubic, Shenzhen, Guangdong, China) was used to demonstrate a multistep capillaric circuit utilizing a 3D-printed SCM. Isopropanol for chip post-processing and Amaranth dye for dried reagent spotting was sourced from Sigma-Aldrich (Burlington, Massachusetts, USA). Colored food dyes (Watkins, Winona, Minnesota, USA) were employed for flow characterization of liquid reagents.

### Proof-of-concept chip fabrication via 3D-printing

We 3D-printed phaseguides and SCMs for feature optimization using a DLP-SLA printer (CADWorks Profluidics 285D) using Clear Microfluidics Resin V7.0a (CADworks3D, Toronto, ON, Canada). Phaseguides, SCMs, and connected microfluidic channels were designed on Solidworks (Waltham, Massachusetts, USA) and exported as Standard Tessellation Language (STL) and sliced to 30 µm layers via the CADworks Utility 6.44 slicer (CADworks3D, Toronto, Ontario, Canada).

The DLP-SLA 3D printer used in our demonstrations has a manufacturer-reported resolution of 20-50 µm in the x- and y-axes but empirical testing shows that consistently high-quality features that match designed dimensions must be closer to 150 µm^10^. Factors affecting this minimum resolution include light diffusion, which can inadvertently cure optically clear resin, and minimization of aliasing due to the printer’s pixel grid. Although smaller features can be achieved with specialized or custom-built printers, our current setup meets the necessary specifications for pinning functionality. Thus, while printing smaller features is technically possible, the additional effort is unnecessary for our current application.

### Capillaric circuit fabrication via low-cost LCD 3D-printing

Chips fabricated for capillaric circuit demonstrations (Fig. 6) were printed using an Anycubic Mono X 6K LCD stereolithography printer (Anycubic Mono X 6K, Phrozen Sonic Mini 8K); a commercially-available, low-cost LCD printer previously characterized by Leong et al^10^. The Chitubox Basic 1.9.5 slicer (Chitubox, Shenzhen, Guangdong, China) was utilized to prepare STL files for printing on the Anycubic Mono X 6K.

### 3D-printing post-processing

After printing, chips printed with CADWorks resin were cleaned with isopropanol (Sigma-Aldrich, Burlington, Massachusetts, USA) and chips printed with Anycubic WaterWash+ were washed with DI water. Chips were dried using a compressed nitrogen gun (Cleanroom World, Centennial, Colorado, USA). This washing and drying was repeated until remaining uncured resin was removed. After drying, the microchips were placed in a Formcure (Formlabs, Somerville, Massachusetts, USA) to post-cure for 1 minute.

Microchips were plasma treated with the Plasma Etch PE-25 (Carson City, Nevada, USA) at 56% power for 1 minute to achieve a hydrophilic surface (θcontact = XX) for capillary-driven flow. Contact angle measurements were made on the Krüss DSA25S Drop Shape Analyzer (Krüss Scientific, Hamburg, Germany). Chips were sealed with ARseal™ 90697 microfluidic tape (Adhesives Research, Glen Rock, Pennsylvania, USA) to seal SCMs and microfluidic channels.

Our routine post-processing is currently dependent on use of microfluidic cover tape. Features are typically printed with open channels, plasma treated for hydrophilicity and then sealed. Tape lacks the rigidity to bridge large negative features like the open reservoirs in phaseguide designs. To address this, we ensured the reservoirs were deep enough to prevent the tape from bowing and sticking to the guides.

### Flow visualization with food dye

Food coloring (1.4% v/v in water) was used to test flow in the capillaric circuits. Nonwoven cleanroom paper (AAWipes, Ann Arbor, Michigan, USA) was used to wick excess liquid off used chips and as a capillary pump in the capillaric circuit demonstration in Fig. 6. Dried amaranth dye spots were prepared by suspending 3% w/w amaranth dye in water and spotting directly onto prepared chips with a custom 3D-printed auxiliary jig to stabilize a pipet tip while dispensing 0.5 µl droplets with a spacing of 2 mm. Once spotted, chips were left to dry for 1 hour. Chips were then sealed with cover tape and tested.

### Image and video capture

Dimensional characterization, image, and video capture were performed on a Keyence VH-S30B digital microscope (Itasca, Illinois, USA).

## Results and Discussion

### Operating principle of phaseguides

Phaseguides are raised rails in microchannels that enable precise control over the liquid filling front. The abrupt height change near the edge of the rail imposes a pressure barrier that guides liquid along the rail before overflowing (Fig.1A.1). Phaseguides do not entirely impede flow and must be shorter than the depth of their housing-reservoir to allow flow to eventually progress. Phaseguide-mediated flow in reservoirs begins with fluid addition to the reservoir inlet, either by forced flow or capillarity. The advancing liquid front fills an area of low pressure until it interfaces with a phaseguide (Fig. 1A.1). A strong Laplace pressure barrier is created by the decreased cross-sectional area of the liquid-air interface and the perpendicularity of the rail, allowing the remaining low-pressure area preceding the phaseguide to continue filling (Fig. 1A.2). Once the area preceding the phaseguide is full and fluid is aligned with the rail, the pressure of the air-liquid interface increases by either forced flow via a pump or capillary pressure on transverse walls that drives flow in the microchannel (Fig. 1A.3). The pinned fluid meniscus stretches, becomes unstable, and breaks at the weakest point along the phaseguide (Fig. 1A.4). In the example shown in Figure 1A, the weakest point along the phaseguide is at the smallest angle between the rail and the side wall. The breakpoint of the phaseguide is tunable and is determined by the inverse relationship between capillary pressure and the interfacing angle, analogous to the relationship between capillary pressure and cross-sectional dimensions in small channels^1,6^. After overflowing at the weakest point, fluid ‘unzips’ over the phaseguide since the Laplace pressure barrier has been overcome (Fig.1A.4). Overflowing fluid fills the next compartment of the reservoir and stops at the next phaseguide, repeating the cycle (Fig. 1A.5).

**Fig.1:**
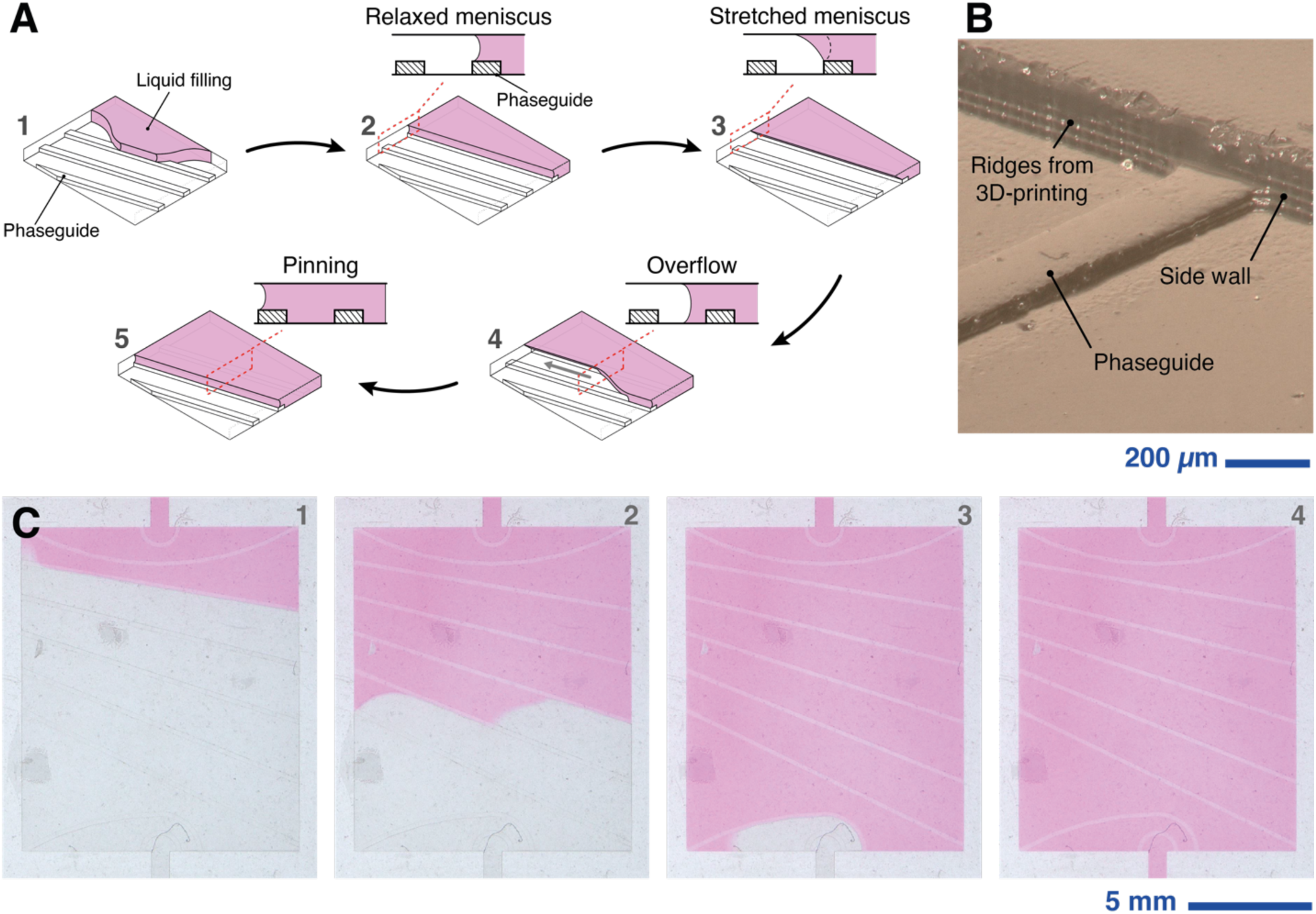
Schematic illustration of phaseguide design and operating principle. (A) Illustration of the control of a liquid filling front using phaseguides in a reservoir. 1. Fluid fills the reservoir and pins at a phaseguide, 2. The meniscus fully aligns with the phaseguide and the meniscus of the air-liquid interface remains momentarily relaxed, 3. The meniscus is stretched by capillary pressure of the fluid interfacing with the device walls, 4. The buildup of capillary pressure is significant enough to breach the phaseguide, causing overflow at the weakest point of the phaseguide, 5. Fluid fills the next segment of the reservoir and pins at the next phaseguide, repeating the cycle. (B) Image of 3D-printed phaseguide (C) Example of bubble-free filling of a reservoir using 3D-printed phaseguides (Movie S1). As laid out in the schematic, **f**luid pins at a phaseguide and with sufficient capillary pressure, overflows. This process cyclically repeats over all phaseguides until the reservoir is filled.

As a proof of concept, we designed and 3D-printed phaseguides that were 200 µm wide and 60 µm tall (Fig.1B). We also showed flexibility of designing and 3D-printing both curved and straight phaseguides. The reservoir was 12 mm x 14 mm x 180 µm for a filled total volume of 29.16 µL. Fluid was effectively pinned by the 3D printed phaseguides and priming behavior was maintained in ordered pinning-and-overflowing processes (Fig.1C, Movie S1). Reservoir filling proceeds via capillary pressure with no bubble formation in the reservoir because of the liquid routing provided by the phaseguides.

### Sequential liquid delivery using phaseguides with distinct pressure barriers

Prior work with cleanroom-fabricated phaseguides established their capability to act as capillary burst valves to allow precise routing of liquids through channel networks^6^. We demonstrated phaseguide-mediated passive valving by designing and 3D-printing rails with different interfacing angles with the channel wall to pre-program distinct pressure barriers (Fig. 2A, Movie S2). We used 3 different phaseguide designs to route liquid between 7 reservoirs in an arrangement that prevented bubble trapping while maintaining the desired flow sequence. All the phaseguides in this demonstration were 3D-printed with the same dimensions (200 µm wide and 60 µm tall), but with different interfacing angles with the side wall: θ_1_ = 30° at all the reservoir inlets, θ_2_ = 90° at the outlet of reservoir 1, and θ_3_ = 150° at the outlets of reservoirs 2 through 7 (Fig 2B).

**Figure 2:**
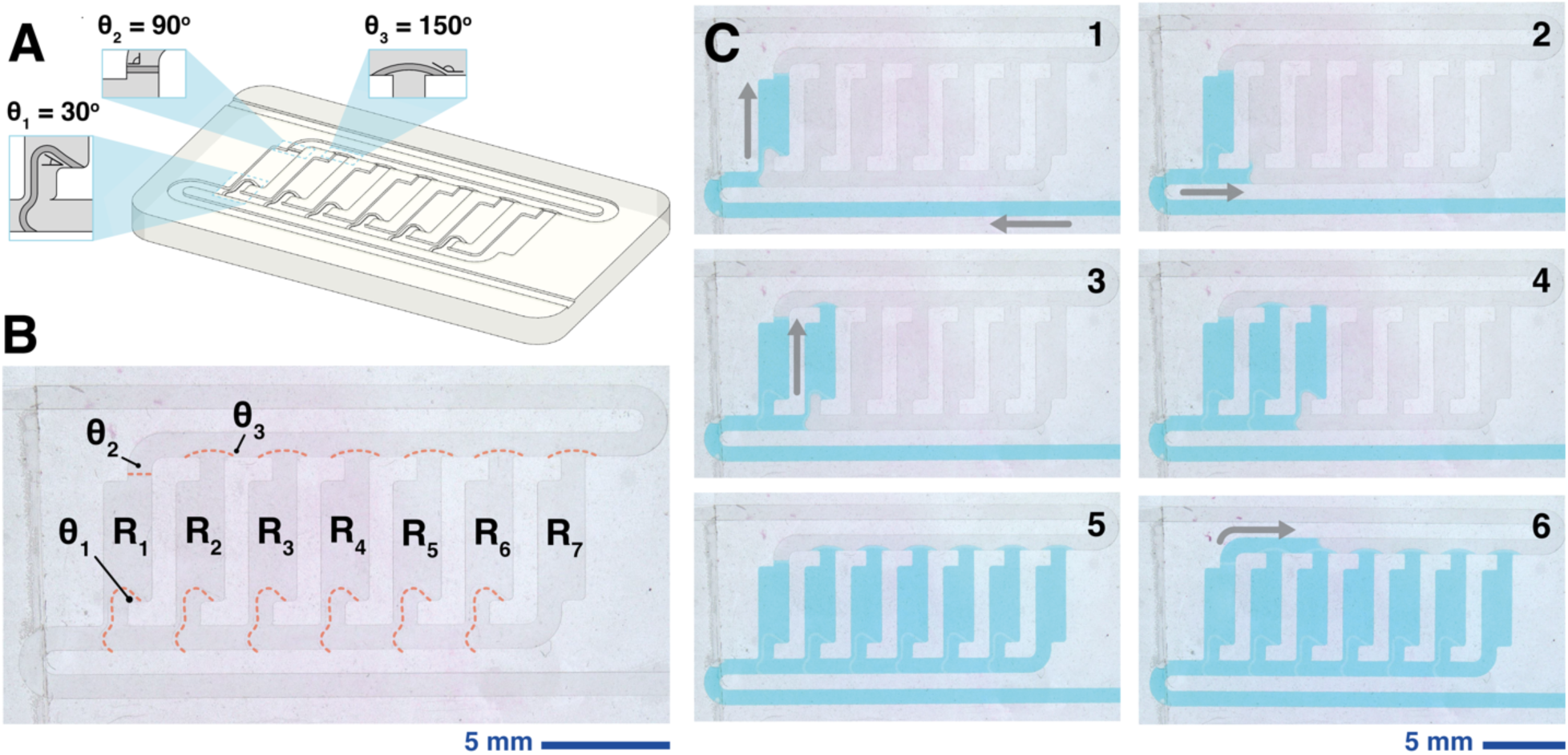
Sequential liquid delivery using phaseguides with distinct pressure barriers. (A) CAD rendering of a sequential valving network of phaseguides where the interfacing angle between the rail and the wall encodes different pressure barriers. The higher the interfacing angle the greater the pressure barrier. (B) Schematic of phaseguides, interfacing angles, and reservoirs labelled on a 3D-printed phaseguide-mediated passive valving network. Interfacing angles, θ_1_ to θ_3_, encode sequential filling of reservoirs from R_1_ to R_7_. (C) Demonstration of a 3D-printed valving network (Movie S2). Phaseguides can sequentially valve reservoirs in a network based on the designed stability of each inlet and outlet^6,34^.

Fluid first enters reservoir 1 and contacts the 30° (θ_1_) phaseguide (Fig 2C.1) which has a pressure barrier that guides flow vertically into the reservoir but not laterally towards the remaining reservoirs. When the fluid reaches the outlet of reservoir 1, it reaches the 90° (θ_2_) phaseguide which has a greater pressure barrier than θ_1._ Liquid is pinned at θ_2_ while overflow occurs at θ_1_ guiding liquid towards the inlet of reservoir 2 (Fig 2C.2). Fluid is once again routed to fill reservoir 2 because of the positioning of phaseguide with θ_1_ near the reservoir inlet. Liquid reaches the outlet of reservoir 2 and contacts a 150° (θ_3_) phaseguide (Fig 2B.3) which has a greater pressure barrier than θ_1_ and θ_2_. Overflow occurs at the phaseguide with the lowest pressure barrier (θ_1_) and liquid again progresses to the next reservoir (Fig 2B.4). This cycle repeats for the reservoirs 3 to 7 (Fig 2B.5). Once all reservoirs are filled, the flow is pinned at phaseguide at θ_2_ at the outlet of reservoir 1 and phaseguide θ_3_ at the outlets of reservoirs 2 – 7. Again, the phaseguide with the lowest pressure barrier (θ_2_) bursts first, and liquid sweeps from the outlet of reservoir 1 through the rest of the channel network to enable bubble free filling of all the reservoirs (Fig 2B.6).

### Key features and operation of self-coalescence modules (SCMs)

SCMs use phaseguides and capillary burst valves to achieve flow redirection or “self-coalescence” that enables uniform reagent rehydration and bioassay automation in microchannels^7,35,36,38^. The phaseguide (leading barrier) in SCMs creates a Laplace pressure barrier that ensures partial reservoir filling until liquid reaches a capillary burst valve (diversion barrier) at the outlet (Fig. 3A). The diversion barrier, aided by the curled end of the leading barrier (Fig. 3C.3), encourages flow redirection that overcomes the Laplace pressure barrier and initiates self-coalescence (Fig. 3C.4) or “unzipping” of flow along the leading barrier. The module unzips until it reaches the end of the SCM body adjacent to the inlet (Fig. 3C.5.), where fluid is pinned by a stop valve at the vent, preventing leakage into the air conduit (Fig. 3B). There are stopping features crucial to SCM functionality that require differentially strong pinning barriers: the leading barrier, the diversion barrier, and the vent barriers. At this point, the SCM body is full, and the diversion barrier becomes unstable via meniscus stretching and capillarity of the device. The diversion barrier spontaneously bursts and the flow proceeds to the outlet (Fig. 3C.6.).

**Figure 3:**
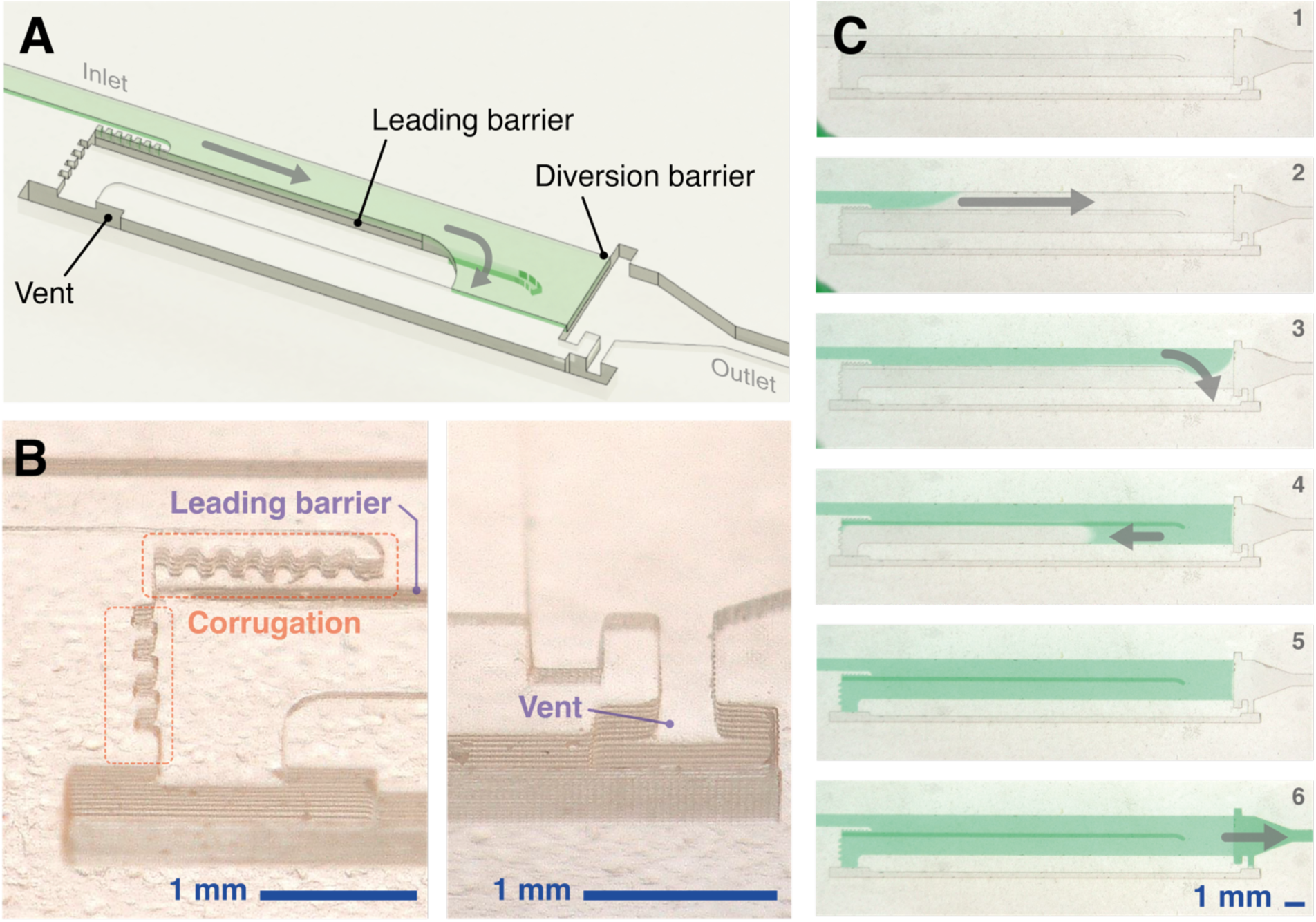
Schematic, key features, and operating principle of 3D-printed self-coalescence modules. (A) Schematic of a typical SCM that labels key features including the leading barrier, the diversion barrier, and vent and illustrates the direction of flow. Flow is guided by the leading barrier, a capillary pinning line, and redirected during filling by the diversion barrier, allowing flow to self-coalesce. (B) Zoomed images of 3D-printed SCMs highlighting additional pinning features added in 3D-printed SCMs for robust fluid pinning. (C) Demonstration of self-coalescing flow in a 3D-printed SCM.

3D-printed SCMs presented here rely solely on capillarity to drive flow so uncontrolled fluid migration is a crucial failure mode that must be prevented. The primary observed failure modes were short circuiting of the self-coalescence by traversing the leading barrier or obstructing reservoir filling by spreading along the back wall and plugging the air vent. Gökçe et al. added corrugations along the leading barrier to prevent short circuiting of flow to the back wall and vent^7^. We included a second set of corrugations at the interface between the back wall and the vent given the larger feature sizes (and consequently, lower pressure barrier) in our 3D-printed devices.

### 3D-printed SCMs scaled up 50X from cleanroom fabrication

We designed SCMs according to the minimum viable dimensions using our benchtop DLP-SLA printer i.e. 30 or 50 µm minimum dimension in the z-direction and 150 µm in the x and y-directions^10^. This required scaling up of the overall device footprint and individual feature sizes, as cleanroom-fabricated SCMs have typical dimensions of 5 – 15 mm long, 0.5 – 1.5 mm wide, and 0.05 mm deep, with volumes ranging from 0.125 – 1.125 µL^7^. As a proof of concept, we designed 3D-printed SCMs that were 19.6 mm long, 2.0 mm wide, and 0.2 mm deep for a total volume of 8.94 µL (Fig.3D). The feature that was scaled up the most in 3D-printed SCMs was the leading barrier, which was 5 µm in cleanroom manufactured devices^7^ and was enlarged to 150 – 200 µm in our 3D-printed devices. The inlet channel and the reservoir/body of the SCM were 200 µm deep. The depths of the leading barrier, diversion barrier, and vent were 500 µm, 400 µm, and 1000 µm, respectively, to encode for different pressure barriers that control the direction of flow in the SCM. Using these dimensions, the 3D-printed SCM exhibits the unzipping or self-coalescing flow (Fig.3D, Movie S3).

3D-printing provides additional degrees of freedom in obtaining multiple channel heights and larger volumes^4,39,40^. We demonstrated that SCM scaling can be applied to accommodate larger liquid volumes, either by proportionally increasing all dimensions of the reservoir (Fig. 4.A.1-2) or by changing only the length (Fig. 4.A.3). We doubled the length of SCM and kept its other dimensions constant to accommodate a 20 µL volume while maintaining self-coalescing flow (Fig.4A, Movie S4). We also showed that we could maintain a compact footprint by adjusting the width and depth of the SCM (increased ∼1.5X each) to achieve the same volume doubling (Fig.4B, Movie S4). We also scaled up the SCM volume to 50 µL, ∼50X the volume of cleanroom-fabricated devices, while still maintaining self-coalescing flow (Fig. 4C, Movie S4). Additionally, we showed that we can arbitrarily change the shape of the capillary pinning line and the overall SCM to fit desired configurations while still maintaining precise control over the liquid filling front, as exemplified by self-coalescence in the W-shaped device (Fig.4D).

**Figure 4:**
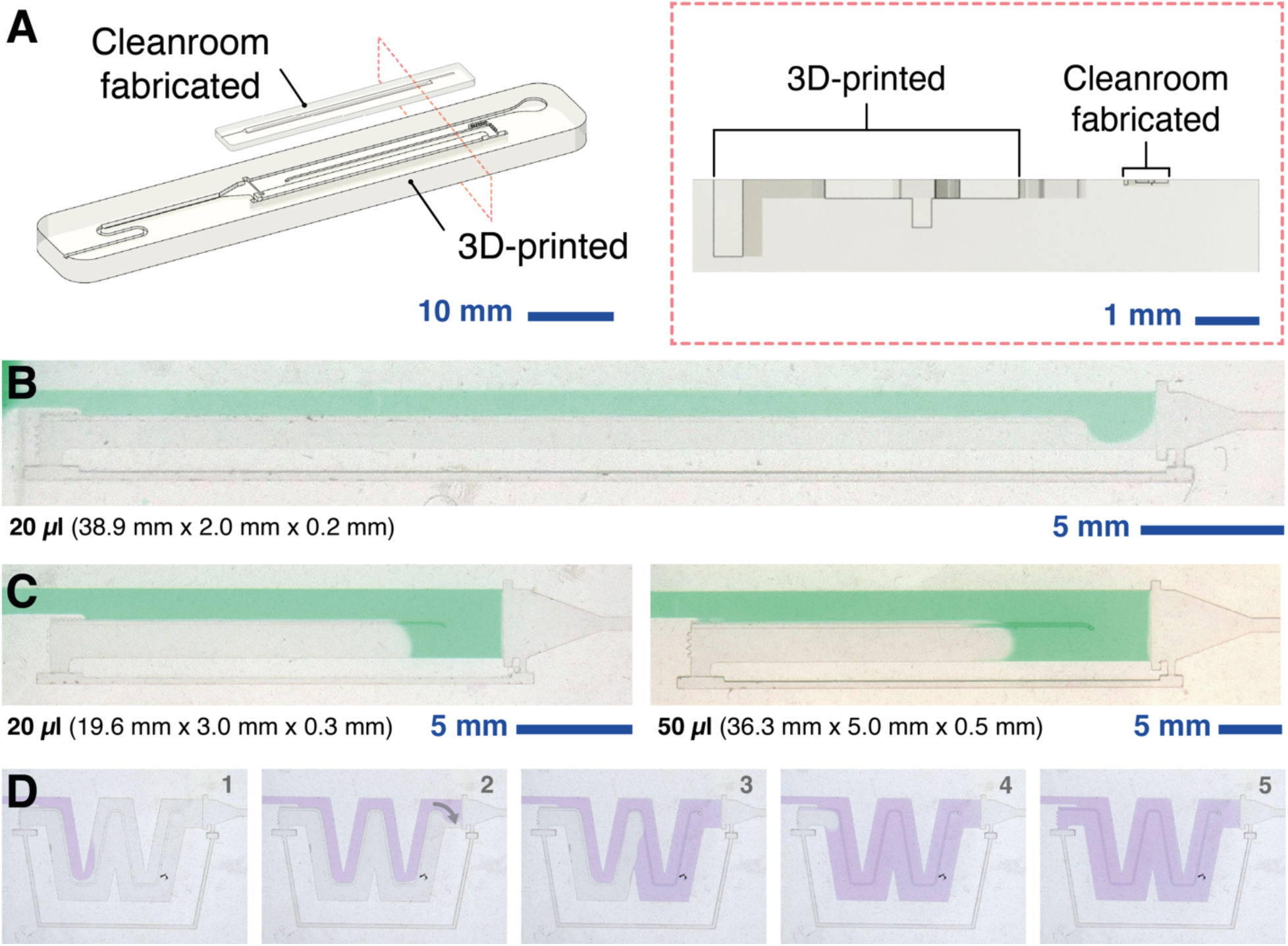
3D printing enables adjustable channel heights and increased volumes in SCMs. (A) Rendering depicting size comparison of traditional cleanroom-fabricated SCM and scaled up 3D-printed SCM. (B) SCM length accommodates a 20 µL volume while preserving self-coalescing flow (Movie S4). (C) Width and depth of SCMs were increased with a fixed ratio to scale to 20 µl and 50 µl (Movie S4). (D) Arbitrarily patterned capillary pinning lines demonstrate precise control over the liquid filling front with an adaptive footprint (Movie S4).

### Proof-of-concept reagent reconstitution using 3D-printed SCMs

Figure 5 demonstrates a proof-of-concept reconstitution of dried reagents in a 3D-printed SCM to visualize pulse shaped dispersion of amaranth spots without accumulation at the liquid front. Using a 3D-printed auxiliary tool that stabilizes a standard pipet tip while dispensing 0.5 µL spots with spacings of 2 mm, we hand-spotted amaranth onto the surface of 3D-printed SCMs. The leading barrier pins fluid as it approaches the diversion barrier (Fig. 5A.1). Once redirected by the diversion barrier, the fluid front self-coalesces, contacting the first dried spot (Fig. 5A.2). Self-coalescence continues, resuspending the dried spots (Fig. 5A.3). Upon complete filling of the SCM, resuspension of the spots did not result in dye accumulation near the diversion barrier (Fig. 5A.4) as would be expected with a reservoir filled without a self-coalescence module^7^. The reconstituted dye continues to diffuse over the course of 40 seconds before the diversion barrier spontaneously bursts. As shown in the cleanroom manufactured demonstrations of SCMs by Gökçe et al., serpentines as fluidic resistors can be added after SCMs to mix and homogenize the reconstituted reagents.

**Figure 5:**
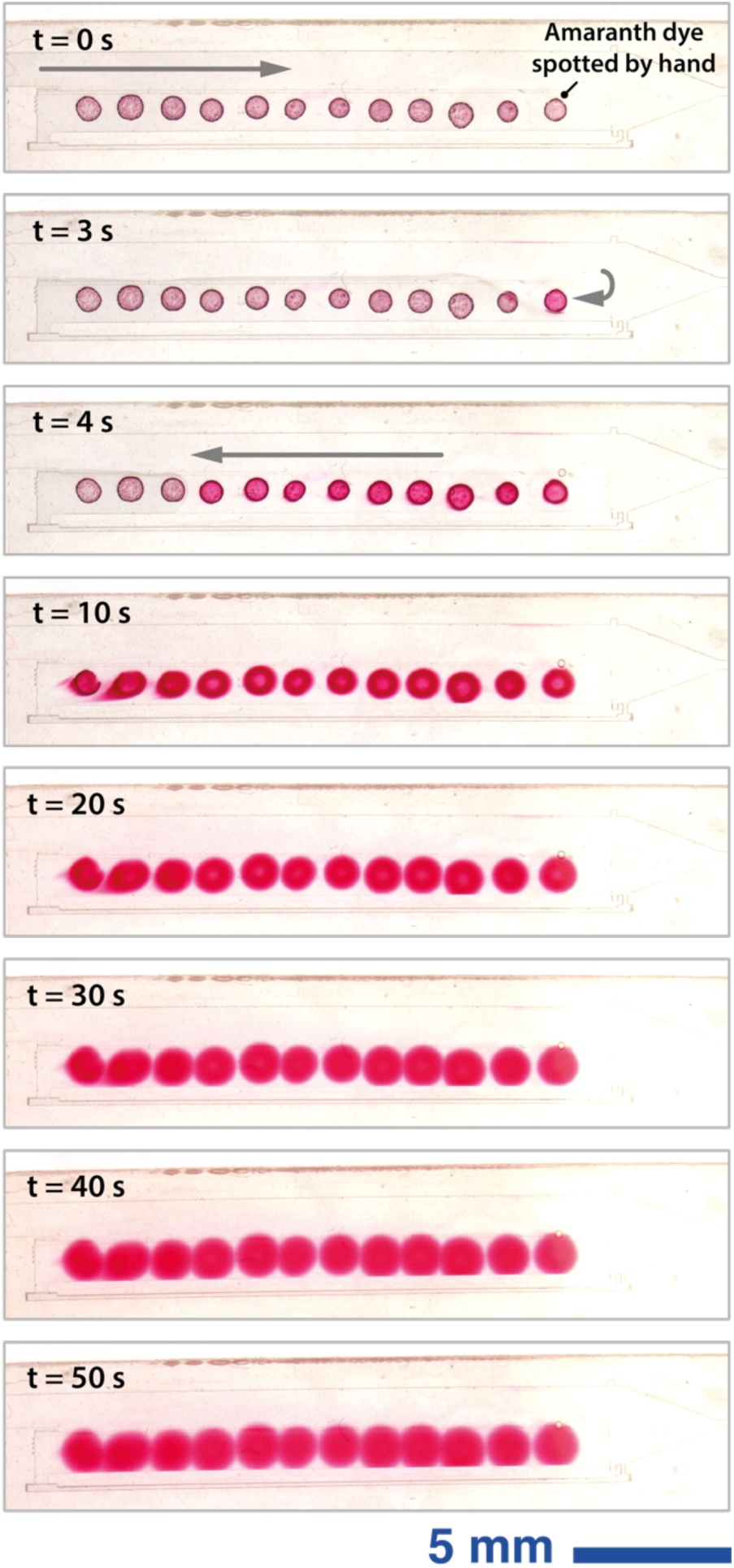
Reconstitution of hand-spotted amaranth dye in large-volume 3D-printed SCM. (A) 0.5 µL spots of amaranth dye spotted on the surface of a 3D-printed SCM with 50 µL volume undergo self-coalescence for resuspension and uniform rehydration. (B) Diffusion of the rehydrated amaranth spots over the course of 40 seconds (Movie S5).

### Integration of an SCM into a 3D-printed capillaric circuit

We demonstrate the feasibility of integrating SCMs into a capillaric circuit with additional flow control elements (Fig.6). This circuit was designed to combine inline reconstitution of a concentrated reagent with dynamic, multistep fluid control. The SCM acts a reservoir that can be preloaded with reagent – in this case a concentrated dye solution – that is terminated by a trigger valve on one end to keep liquid in place until a liquid control signal is provided, and a retention burst valve on the other end to enable pre-programmed sequential drainage (Fig.6A).

**Figure 6:**
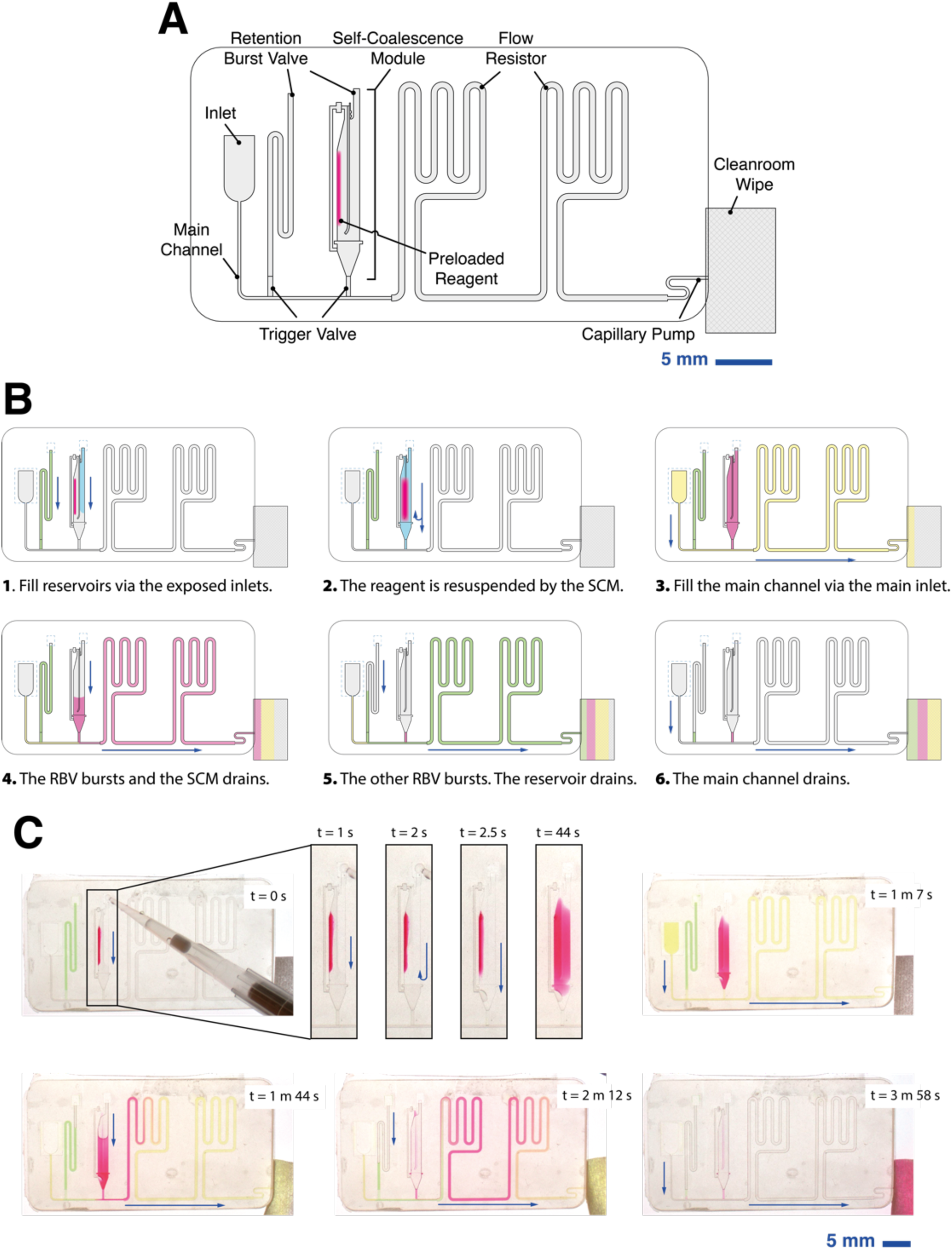
Integration of an SCM into a low-cost 3D-printed capillaric circuit. (A) Schematic of a capillaric circuit that includes an SCM, trigger valves, retention burst valves, flow resistors and a capillary pump. The SCM acts a reservoir that can be preloaded with reagent – in this case a concentrated dye solution – that is terminated by a trigger valve on one end to keep liquid in place until a liquid control signal is provided, and a retention burst valve on the other end to enable pre-programmed sequential drainage. The circuit supports inline reagent reconstitution and pre-programmed, instrument-free, multi-step fluid control. (B) Schematic illustrating expected operation of capillaric circuit with integrated SCM. (C) Timelapse images showing observed pre-programmed delivery with capillaric circuit containing SCM (Movie S6).

To visualize suspension of a concentrated reagent in the SCM, we pipetted 1 µL of food dye onto the surface of the chip. In further applications, reagents could be spotted onto the chip surface using an auxiliary support as in Fig. 5 or by a small volume deposition spotter. However, resuspending a concentrated liquid food dye demonstrates the flexibility to incorporate SCMs as a tool for effective dilution of liquid reagents or samples in future applications.

Self-coalescence aided dilution of a concentrated liquid happens during loading of the reagent reservoirs (Fig. 6B.3-4). Once reservoirs are filled, fluid is pinned at trigger valves until actuation by the addition of liquid to the inlet of the main channel (Fig. 6B.5). The weakest RBV sits atop the SCM and bursts when liquid is pinned at the RBV at the main channel inlet (Fig. 6B.6). The “tug of war” style bursting mechanism of RBVs means that we can design the geometry of the inlet of the SCM so that it bursts first while the main channel inlet and other reservoir of the capillaric circuit hold their liquid in place. The patterning of the capillary pinning lines within the SCM do not compromise rservoir drainage. When the SCM reservoir is completely empty, the second RBV (with the second lowest capillary pressure in the circuit) bursts (Fig. 6B.7). Finally, the main channel drains, completing the circuit.

This integrated capillaric circuit with an SCM was fabricated using a low-cost (Anycubic Mono X 6K, MSRP ∼$340 USD as of December 19, 2024) LCD-SLA printer using inexpensive WaterWash+ resin (Anycubic, MSRP $29 USD/kg). The printer is ∼50X cheaper than the DLP-SLA printer used for initial characterization of SCMs and demonstrates the potential for rapid prototyping of SCMs with minimal capital and operating costs^10^.

### Comparisons with other published work

To our knowledge, examples of 3D-printed phaseguides in the literature have so far been limited to two-photon polymerization, which is expensive and still requires cleanroom facilities^41^, or scalloped channel edges in hydrogels that exhibit phase guiding behavior^40^. There was also a recent demonstration of laser cutting of capillary pinning lines in microchannels to rehydrate reagents^37^. Although that approach was inspired by SCMs and shows the capability for rapid and inexpensive prototyping of capillary pinning lines, key fluidic elements like vents and diversion barriers were not incorporated, thereby precluding the ability to showcase the full functionality and integration of SCMs within capillaric circuits. There have also been calls for vat polymerization of 3D-printed phaseguide-mediated organ-on-a-chip devices, but no demonstrations have been presented thus far in the field^39^.

In comparison, our work shows functional 3D-printed phaseguides despite the greater surface roughness and layer-by-layer fabrication process of benchtop 3D printing compared with conventional cleanroom fabrication. We also 3D-printed functional SCMs and demonstrated that they can be scaled by applying a universal scaling factor to the height, width, and length of the reservoir body (Fig. 4.A.1-2) or by extending only the length while keeping height and width fixed (Fig. 4.A.3). In line with findings by Gökçe et al., shallow, wide SCMs exhibit more stable aspect ratios due to minimized liquid-air interface at the leading barrier, suggesting that maintaining an optimized aspect ratio is beneficial during scaling^7^. Lengthening the SCM provides a straightforward scaling method, but depending on the device layout, this can result in long footprints and high fluidic resistance, which may downstream processes/applications. In testing a 100 µL SCM, the largest volume we designed, we encountered difficulties in uniform filling upon redirection by the leading barrier, underlining the importance of asymmetrical CPL placement relative to the SCM body’s centerline for large volumes. Gökçe et al.’s work shows that asymmetrical CPLs do not hinder self-coalescence and can accommodate specific spotting patterns, suggesting this approach could be valuable in future work if larger SCM volumes become necessary.

### Exceptions and unaddressed aspects of 3D-printed phaseguides and SCMs

We optimized the design of phaseguides and SCMs mainly by empirical observations given the rapid prototyping offered by 3D-printing. Further work could be done to integrate theoretical models of the air-liquid interface^7^ at phaseguides and also characterize the impact of 3D-printing fabrication imperfections like feature sharpness and perpendicularity.

Although we showed sequential filling of 7 reservoirs using 3 different phaseguide strengths, we have not yet explored the limit of discretized steps that could be automated by 3D-printed phaseguides. We anticipate that, similar to past work with capillary retention burst valves, the resolution and precision of 3D-printing will limit the number of distinct pressure barriers that can be geometrically encoded^4^. We also expect that the imperfections in the perpendicularity of phaseguides in the z-axis introduced by the layer-by-layer nature of stereolithographic 3D printing could introduce weaknesses or nonlinearities in the pressure barriers, affecting their stability.

One major unaddressed aspect of our work is that we did not incorporate precision-deposited dried reagents into the 3D-printed SCMs. Instead, our work focused on the feasibility of fabricating functional phaseguides and SCMs using inexpensive benchtop 3D-printing. Although we observed the expected unzipping of flow and lack of accumulation of rehydrated reagents at the fluid front in Fig. 4, the imprecision of our manual spotting method limited the ability to quantitatively evaluate reagent rehydration in 3D-printed SCMs to the same extent as was reported with cleanroom-manufactured devices. Future work is required to validate 3D-printed SCMs with dried reagents. This could require exploring proven strategies for enhancing lateral reagent homogenization in cleanroom-fabricated SCMs such as flow mixers at the outlet of SCMs^7^. Additionally, self-coalescence across differential segments of SCMs—curved versus straight— requires validation to ensure effective reagent reconstitution and optimization of spot placement for specific assay applications.

This unaddressed aspect of validation highlights current limitations to democratization of this technology. Despite significant advances in rapid prototyping via low-cost 3D printers, reliance on expensive (and often centralized) small volume reagent deposition spotters still limits experimentation and application of this technology to highly resourced research settings. Incorporation of frugal reagent deposition systems, like modified office inkjet printers^42^, could help increase the adoption of SCMs without high capital costs.

Broad use of SCMs and integration with other capillaric fluidic elements and circuits could offer many advantages in the development of point-of-care diagnostics. Dried reagents that do not require cold chain storage can reduce the number of liquid loading steps, enable long-term storage and distributed deployment, and improve the user-friendliness of microfluidic devices^7^.

## Conclusion

We 3D-printed phaseguides and SCMs and showed comparable performance to cleanroom-fabricated features. We showed that 3D-printed phaseguides could guide the liquid filling front without bubble formation in reservoirs. Using phaseguides with different strengths, preprogrammed by the angle between the phaseguide and the channel walls, we also demonstrated sequential filling of multiple reservoirs within the same 3D-printed device. We also used 3D-printed phaseguides as pinning features to obtain the characteristic self-coalescence or unzipping flow in SCMs. 3D-printing allows us to rapidly manufacture SCMs, easily scaling them to feature sizes up to 50X larger than those of cleanroom-fabricated devices. We also showed that we could change the shape of the 3D-printed SCMs to reconfigure device footprint and/or volume. Finally, we integrated 3D-printed SCMs with other capillaric elements – including trigger valves, retention burst valves, flow resistors, and capillary pumps – to demonstrate dilution of pre-loaded liquid and pre-programmed, instrument-free, multi-step liquid delivery.

The 3D printing and integration of SCMs into capillaric circuits represents a significant advancement, combining rapid and inexpensive manufacturing with robust and scalable fluid control. While cleanroom fabrication has traditionally been essential for developing microfluidic control elements, especially features like phaseguides and SCMs that precisely manipulate capillary pinning lines, 3D printing has expanded both accessibility and versatility, enabling cost-effective prototyping without compromising functionality. This work broadens the range of fluidic elements that can be quickly produced, facilitating applications in assay automation, organ-on-chip systems, and diagnostics by addressing previous limitations in fluid control and reagent reconstitution. To further extend the utility of these systems, future designs could network phaseguides with additional capillaric elements, eliminating the need for external pumps and opening new possibilities in point-of-care diagnostics. Altogether, the integration of 3D-printed capillaric components marks a critical step in democratizing liquid handling technology to enable broader, impactful applications by scientists in high- and low-resource settings.

## Acknowledgements

We would like to thank Dr. Ashleigh Theberge for her valuable feedback and motivation to share our findings with the broader scientific community. We would also like to thank Dr. Lucas Meza and the Meza Lab for access to instruments. We are grateful for funding from the Arnold and Mabel Beckman Foundation.

## Notes

### Competing Interest Statement

The authors have declared no competing interest.

